# Architecture of human interphase chromosome determines the spatiotemporal dynamics of chromatin loci

**DOI:** 10.1101/223669

**Authors:** Lei Liu, Guang Shi, D. Thirumalai, Changbong Hyeon

**Author notes:** L.L., G. S., D. T, and C.H. designed and performed research, analyzed data, and wrote the paper. The authors declare no conflict of interest.

## Abstract

By incorporating the information of human chromosome inferred from Hi-C experiments into a heteropolymer model of chromatin chain, we generate a conformational ensemble to investigate its spatiotemporal dynamics. The heterogeneous loci interactions result in hierarchical organization of chromatin chain, which obeys compact space-filling (SF) statistics at intermediate length scale. Remarkably, the higher order architecture of the chromatin, characterized by the single universal Flory exponent (*ν* = 1/3) for condensed homopolymers, provides quantitative account of the dynamical properties of the chromosome. The local chromosome structures, exemplified by topologically associated domains (~ 0.1 − 1 Mb), display dynamics with fast relaxation time (≲ 50 sec), whereas the long-range spatial reorganization of the entire chromatin 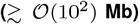 occurs on a much longer time scale (≳ hour), suggestive of glass-like behavior. This key finding provides the dynamic basis of cell-to-cell variability. Active forces, modeled using stronger isotropic white noises, accelerate the relaxation dynamics of chromatin domain described by the low frequency modes. Surprisingly, they do not significantly change the local scale dynamics from those under passive condition. By linking the spatiotemporal dynamics of chromosome with its organization, our study highlights the importance of physical constraints in chromosome architecture on the sluggish dynamics.

**Significance Statement:** Chromosomes are giant chain molecules made of hundreds of megabase-long DNA intercalated with proteins. Structure and dynamics of interphase chromatin in space and time hold the key to understanding the cell type-dependent gene regulation. In this study, we establish that the crumpled and space-filling organization of chromatin fiber in the chromosome territory, characterized by a single universal exponent used to describe polymer sizes, is sufficient to explain the complex spatiotemporal hierarchy in chromatin dynamics as well as the subdiffusive motion of the chromatin loci. While seemingly a daunting problem at a first glance, our study shows that relatively simple principles, rooted in polymer physics, can be used to grasp the essence of dynamical properties of the interphase chromatin.

---

The organization of chromosomes, comprised of a long DNA/chromatin chains, depends on the length scale. On ≲ 10 nm scale, dsDNA wraps around histone octamers to form nucleosomes, which assembly constitutes the chromatin fiber. The signature that the chromatin fiber is further compacted into higher-order structures, such as topologically associated domains (TADs) and chromatin compartments has come from the interaction patterns inferred from Hi-C data (1–3).

The three dimensional (3D) structures of chromatin vary with the developmental stage (4) and cell types, which has resulted in the appreciation that chromatin structure plays in the regulatory role. For long range transcriptional regulation (5–7), two distal genomic loci has to be in proximity. Hi-C maps of chromatin, measuring mean contact frequencies between cross-linked DNA segments from an ensemble of millions of fixed cells, suggest its hierarchical organization. Chromosomes at ~ 5 Mb resolution are partitioned into alternating A and B type compartments that are enriched with active and inactive loci, respectively (1). At a higher resolution the data reveals the formation of TADs, the submegabase sized functional building blocks of interphase chromosome (2). While the chromatin chain within TADs is highly dynamic (8), the boundaries between the TADs are well insulated across different cell types. Genome-wide Hi-C maps at even higher resolutions of 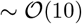 Kb indicate at least 6 subcompartment types, characterized by distinctive histone markers and chromatin loops (3). In addition fluorescence images give glimpses of real-time chromatin dynamics *in vivo* (9–12), allowing us to decipher the link between structure, dynamics, and function (13–15).

Advances towards the mechanistic underpinnings of chromatin compaction are also being made using theory and computations. Based on the knowledge of the convergent orientation of the CTCF-binding motifs, the loop extrusion polymer model (16, 17) was proposed to explain the formation of TADs and predict the contact maps of edited genomes upon deletion of CTCF-binding sites (16, 17). While homopolymer models with geometrical constraints (1, 18–22) capture the physical basis of chromosome organization, one can utilize the information from Hi-C and fluorescence *in situ* hybridization to sharpen the model (23–26).

To explore the spatiotemporal dynamics of a chromosome, we modified a recently developed heteropolymer model, – Minimal Chromatin Model (MiChroM) – whose parameters were trained based on the Hi-C data of Chr10 (chromosome 10) from human B-lymphoblastoid cell (27). The ensemble of chromosome structures generated from the MiChroM could faithfully reproduced the experimental Hi-C maps of all other autosomes (27). The resulting chromosome structures were characterized with the paucity of entanglements, phase separation of A/B compartments, and enrichment of open chromatin chain at the periphery of chromosome territories (27). Furthermore, the multistability of free energy landscape for individual chromosomes (28) rationalizes the cell-to-cell variability observed in single-cell Hi-C data (7, 29, 30).

The primary aim of this study is to elucidate the physical principles underlying the chromatin dynamics, which has received much less attention. To this end, we imposed the chain non-crossing constraint on the chromosome structures generated from MiChroM, and carried out Brownian dynamics simulations. Our study shows that the basic features of chromatin dynamics observed in experiments is intimately connected to the crumpled, hierarchical, and territorial organization of interphase chromosomes. By incorporating active forces onto *active* loci, we address the extent to which the activity contributes to the dynamic properties of the interphase chromatin.

## Results and Discussions

### Heteropolymer model for chromosome

In MiChroM (27), each chain monomer represents 50 Kb of DNA segment. As a consequence the model describes chromosome organization on large length scale, a feature that is crucial for dynamics. Based on the correlation between the distinct patterns of inter-chromosomal contacts and epigenetic information, MiChroM assigns one of six types of subcompartments (B3, B2, B1, NA, A1, and A2) to each CG monomer (3). In the Hi-C map, candidate binding sites for CTCF (16) or lamin A (12) show much higher contact frequencies than their local background. As a result, the interactions between monomers are accounted for by the potential of a homopolymer, monomer type dependent interactions, attractions between loop sites, and genomic distance-dependent condensation energies (See Supporting Information (SI)). We confine the chromatin to a sphere with a volume fraction of 10 %.

To sample the chromatin conformations at equilibrium, we performed Langevin simulations at low friction (31) (see SI Text). The resulting conformational ensemble of Chr10 captures the “checkerboard” pattern of the Hi-C contact map (3) (Fig. 1A), and reproduces the characteristic scaling of contact probability *P*(*s*) ~ *s*^−1^ over the intermediate range of genomic distance 1 < *s* < 10 Mb (Fig. S1B). The distribution of Alexander polynomial, |Δ(*t* = –1)| (32)(Fig. S1D), characterizing chain entanglement, has the highest mode at zero, indicating that the majority of chromosome conformations are free of knots. The radial distributions of monomers belonging to the different type of subcompartment (27, 33) reveal that in contrast to the condensed and transcriptionally inactive loci, which are buried inside the chromosome, open and active loci are enriched near the surface, which presumably improves the accessibility of transcription factors (Figs. S1E, S1F).

**Fig. 1.**
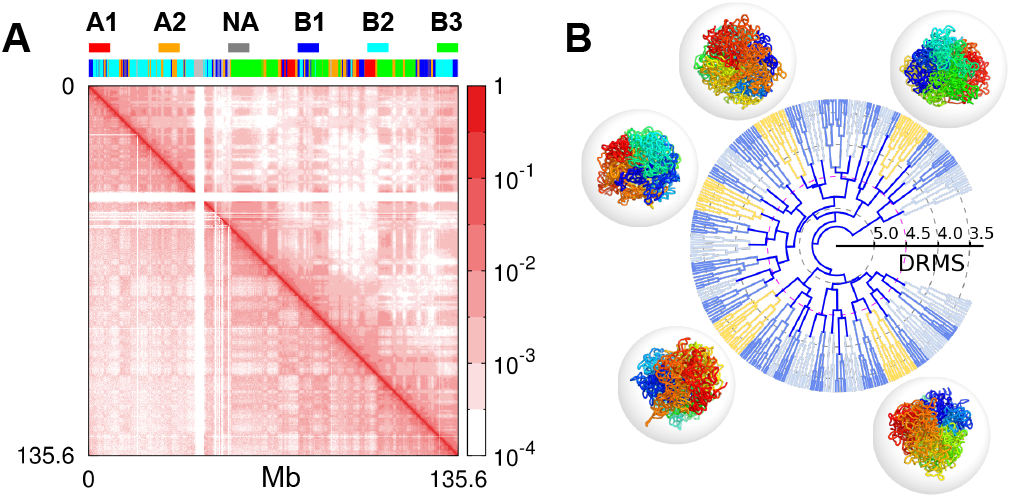
Conformational ensemble of Chromosome 10 of human B-lymphoblastoid cells generated from simulations. (A) The contact frequency map from the ensemble of structures generated using MiChroM (upper right corner) reproduces the overall checkerboard pattern of Hi-C map (lower left corner). The 6 subcompartment types assigned to chromosome loci are depicted on the top. (B) The dendrogram represents the outcome of hierarchical clustering of the heterogeneous ensemble of structures. The distance between leaf nodes (structures) *k* and *l* is given by 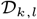 (Eq.1), and the distance between two clusters *K* and *L* is defined as 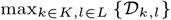. Among the clusters whose inter-cluster distance is smaller than 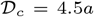, the centroid structures (*k*_c_ ∈ *K*), which minimize 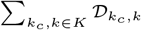, are displayed in rainbow coloring scheme for the 5 most populated clusters (colored in yellow).

Substantial heterogeneity of structures is identified in the conformational ensemble. We use the distance-based root-mean-square deviation (DRMS, 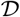),

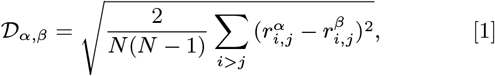

to quantify the similarity between two conformations and partition the conformational ensemble into multiple clusters. In this clustering method, two chromosome structures, say *α* and *β*, that are within a cut-off value 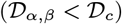 are considered similar and grouped together. We carried out hierarchical clustering by repeating this procedure by varying the value of 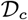 to produce a dendrogram (Fig. 1B); the ensemble is decomposed into many clusters (see Fig. S2). At 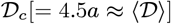, distinction between the structures belonging to different clusters is visually clear (Fig. 1B), suggesting the cell-to-cell variability seen in the recent single-cell Hi-C data (7, 29, 30). The partitioning of the conformational ensemble into distinct cluster is a first indication that the folded landscape of chromosome is rugged. Consequently, we expect that the underlying dynamics should exhibit glass-like behavior (22).

### Subdiffusive dynamics of chromatin loci

An ensemble- and time-averaged mean square displacement (MSD) for chromatin loci was calculated by analyzing the trajectories. The time averaged MSD of a loci is 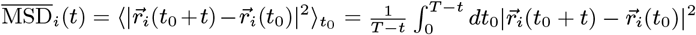, and the ensemble averaged MSD is obtained using 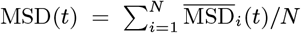. As shown by the MSD curve in Fig. 2A, the diffusion of chromatin loci is characterized by three different time regimes. For a very short time interval (*t* < 10^-2^*τ*_BD_), the loci diffuse freely with MSD ~ t. At the intermediate time scale, corresponding to the Brownian time *t* ~ *τ*_BD_ ~ *α*^2^/*D*, each monomer starts to feel the neighboring monomers along the chain. For *t* > 10^3^ *τ*_BD_, a subdiffusive behavior of MSD ~ *t^β^* with *β* ≈ 0.4 is observed. This exponent is in line with the reported values of *β* = 0.38 ~ 0.44 (34) and *β* = 0.4 ~ 0.7 (12) in live human cells, and is also in reasonable agreement with the diffusion exponent *β* = 0.32 ± 0.03 measured for the whole genome of ATP-depleted HeLa cells (9).

**Fig. 2.**
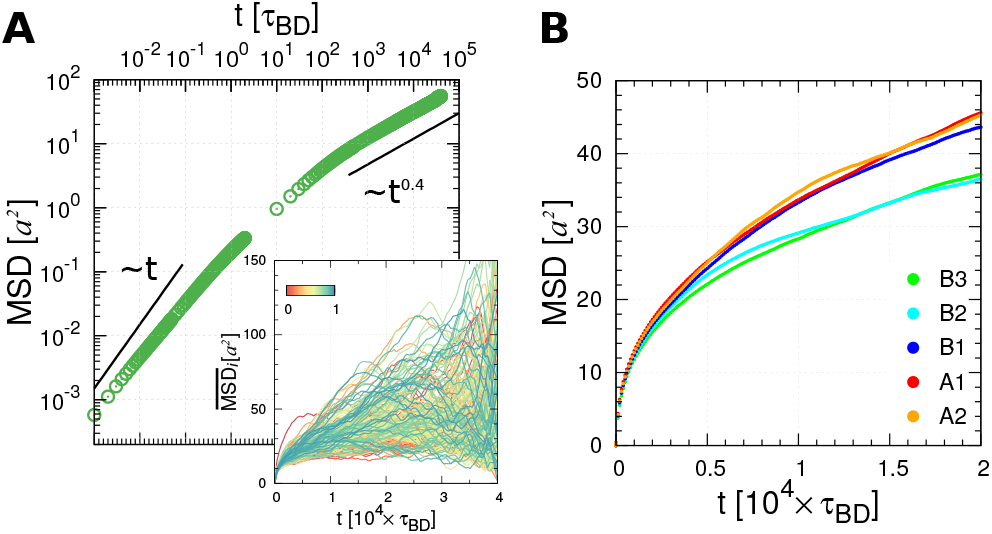
Subdiffusive behavior of chromatin loci. (A) Ensemble- and time-averaged MSD of loci with time in a log-log plot (Inset displays the time-averaged MSD for individual loci, color-coded by a normalized monomer index *i/N*). (B) MSD of loci with different subcompartment types (3)

The exponent *β* = 0.4 can be rationalized using the following argument. The spatial distance (*R*) between two loci separated by the genomic distance, *s*, satisfies *R*(*s*) ~ *s*^*ν*^, where *ν*, the Flory exponent (20, 35), is *ν* = 1/2 for ideal chain obeying Gaussian statistics, and *ν* = 1/3 for space-filling (SF) chain. Notice that the MSD of an expanded locus of arc length s scales with time *t* as MSD ~ *t^β^* ~ *D*(*s*) × *t* ~ *D_o_* × *t*/*s*, where the scaling relationship of diffusion constant of freely draining chain *D*(*s*) ~ *D_o_*/*s* is used. The use. of the relation of MSD ~ *R^2^* (*s*) ~ *s*^2*ν*^ allows us to relate *s* with *t* as *s* ~ *t*^*β*/2*ν*^. It follows that MSD ~ *t^β^* ~ *t*^1−*β*/2*ν*^, giving *β* = 2*ν*/(2*ν* + 1) (34). Thus, we obtain

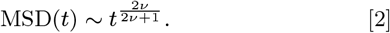

The SF organization of chromosome implies *ν* = 1/3, and hence *β* = 0.4, which explains our BD simulation result at *t*/*τ*_BD_ ≫ 1. A similar argument was used to explain the time-dependence of MSD(*t*) found in an entirely different model of chromosomes (36).

Meanwhile, it has recently been shown using high-throughput chromatin motion tracking in living yeast that MSD~ *t*^0.5^ for all chromosomes (10). The yeast chromosomes obey Gaussian statistics, *R*(*s*) ~ *s*^1/2^ and *P*(*s*) ~ *s*^−3/2^, indicative of *ν* = 1/2. Evidently, from Eq.2, MSD~ *t*^1/2^ (10). Therefore, Eq.2 suggests that the diffusivity of loci is closely linked to the global architecture of chromatins (34, 37).

### Euchromatin versus heterochromatin dynamics

According to a recent single nucleosome imaging experiment (34), diffusion of the heterochromatin-rich loci in the nuclear periphery is slower than the euchromatin-rich loci in the interior. The time-averaged MSD (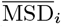) exhibits substantial dispersion among different loci (Fig. 2A, inset). Depending on the subcompartment types, loci move with different diffusivity (see Fig. 2b). The A-type loci, which are less condensed and close to the chromosome surfaces, diffuse faster than the loci of type B2 and B3. Similarly, transcriptionally active loci move slightly faster compared with inactive ones. Although the diffusivity is greater for the active loci, they still have the same *β* = 0.4, which suggests that the chain architecture is the key determinant of the diffusion exponent. Below we will show that even if active forces are incorporated into the dynamics, the value of *β* is unchanged.

### Correlated loci motion in space and time

For complex systems like genomes or chromosomes, various correlation functions can be used to quantify the dynamic properties of the system in space and time.

#### Correlation in time

We first calculated the correlation function of displacement of the *i*^th^ and *j*^th^ loci divided by waiting time Δ*t*, which defines the mean velocity correlation function (11, 38),

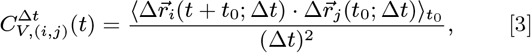

where 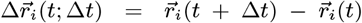 and 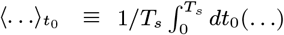, with *T_s_* being the total simulation time, denotes an average over time to. Regardless of Δt, the autocorrelation function 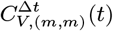 calculated for the midpoint monomer (*m* = *N*/2) displays a negative correlation peak 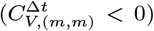 at *t* = Δ*t* (Fig. 3a), followed by a slow relaxation to 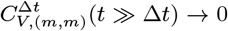. The curves plotted with the rescaled time *t*/Δ*t* nicely overlap onto each other, thus allowing us to assess the variation among the curves (Fig. 3b).

**Fig. 3.**
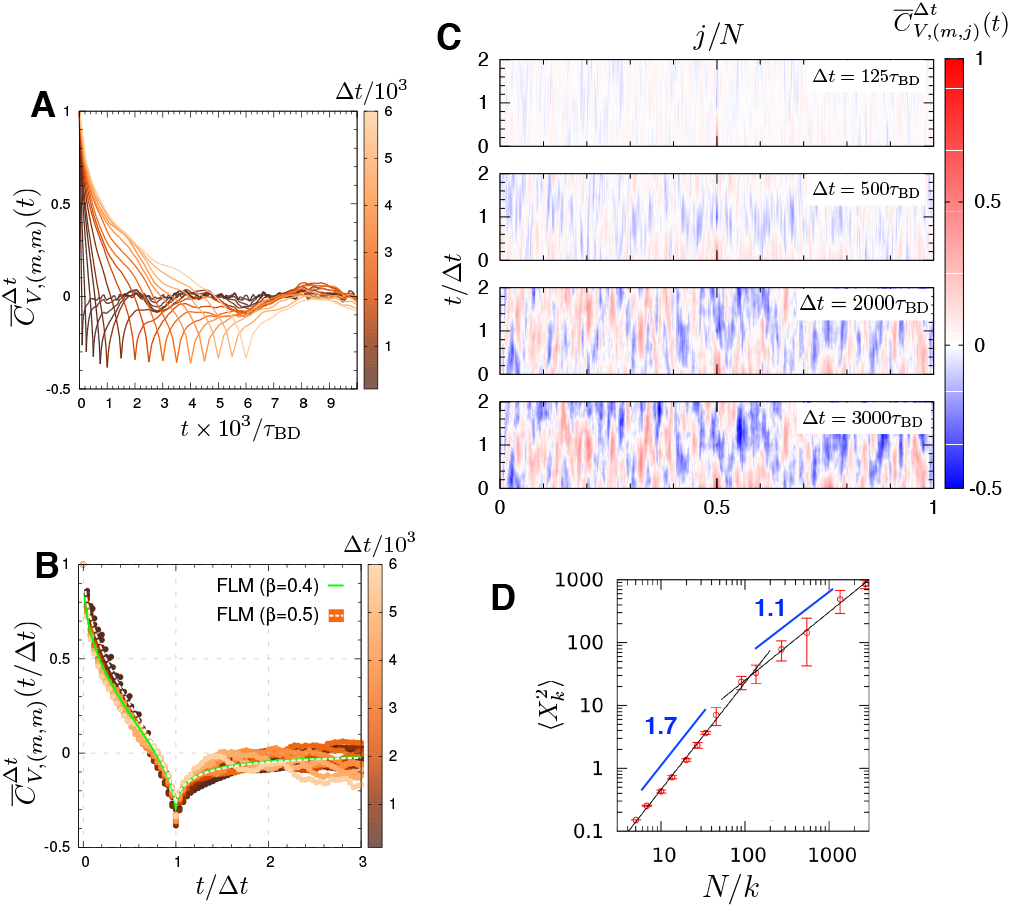
Temporal correlation of locus dynamics calculated from the simulations. 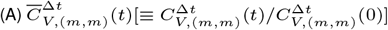 is a normalized mean velocity autocorrelation calculated for the midpoint monomer. Different lines represent different lag times, from Δ*t* = 100 *τ*_BD_ (dark) to 6000 *τ*_BD_ (light). (B) The correlation functions with rescaled argument, 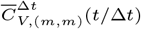. The theoretical curves calculated by assuming the fractional Langevin motion (38, 39) are plotted. The theoretical curve are: 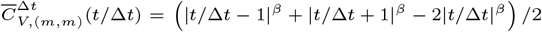 with *β* = 0.4 (green) and *β* = 0.5 (white dashed line for the Rouse chain). (C) Mean velocity cross-correlation between the midpoint (*i* = *m* = *N*/2) and others (*j*), 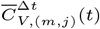 for increasing lag time Δ*t* = 200, 500, 2000, 3000 *τ*_BD_ from the top to bottom. (D) Scaling relation of Fourier modes *X_k_* with 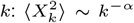 with *α* = 1.7 for large *k* and *α* = 1.1 for small *k*.

Based on the interpretation of fractional Langevin motion, one could posit that the dynamic behavior of chromatin locus captured in 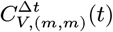 is caused by viscoelasticity of the effective medium (40). However, even the ideal Rouse chain in free space (β = 0.5) displays a similar curve 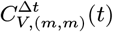 (Fig. 3B). For the Rouse chain in free space, the negative correlation peak, which arises from restoring forces acting on the monomer, is solely due to the chain connectivity with the neighboring monomer along the chain. As Fig.3B shows that the difference between 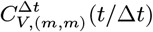 with *β* = 0.4 for the chromatin model and with *β* = 0.5 for the Rouse chain is subtle, and not easy to discern.

In theory, the behavior of our chromatin model can be distinguished from the Rouse chain by calculating the Fourier modes, 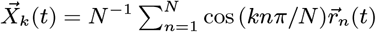. While 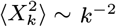 is anticipated for the free Rouse chain (41), we find 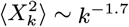 for large *k* values (*N/k* ≲ 100. See Fig. 3D). The Fourier modes for chromatin are expected to scale 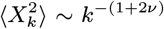 (34). Thus, the exponent of 1.7 is explained again by the SF statistics with *ν* = 1/3.

Cross-correlations of mean velocity between the midpoint (*i* = *N*/2) and other loci (*j* ≠ *N*/2) show how the correlation of our chromatin model changes with time (Fig.3C). In contrast to the viscoelastic Rouse polymer model (39), the mean velocity cross-correlation reveals non-uniform and undiminishing correlation pattern, which suggests that the chromosome structure is maintained through heterogeneous loci interactions defying complete equilibration, an indication of glassy dynamics.

#### Correlation in space

Recently, displacement correlation spectroscopy (DCS) using fluorescence, employed to study the dynamics of a single nucleus, has revealed that a coherent motion of the *μ*m-sized chromosome territories could persist for *μ*s to tens of seconds (9). In order to provide structural insights into these findings, we studied the spatial correlation of the chromosome structure. The spatial correlation between chromatin loci from our simulations can be evaluated using

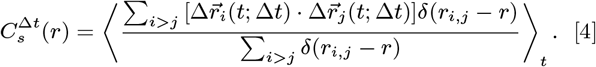

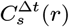 quantifies the displacement correlations between loci separated by the distance *r* over the time interval Δ*t*. 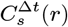 decays more slowly with increasing Δ*t*. The correlation length calculated using 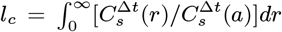, shows how *l_c_* increases with Δ*t* (Fig.4B). To paint an image of displacement correlation over the structure, we project displacements of the monomers near the equator of the confining sphere (−*a* ≤ *z* ≤ *a*) onto the *xy* plane, and visualize the dynamically correlated loci moving parallel to each other by using similar colors (see Fig. 4C). If Δ*t* < 100 *τ*_BD_, the spatial correlation of loci dynamics is short-ranged and the displacement vectors appear to be random. But, with a longer waiting time (Δt > 500 *τ*_BD_), we observe multiple groups of coherently moving loci that form substantially large domains (~ 5*a* ≈ 0.75 *μm*).

**Fig. 4.**
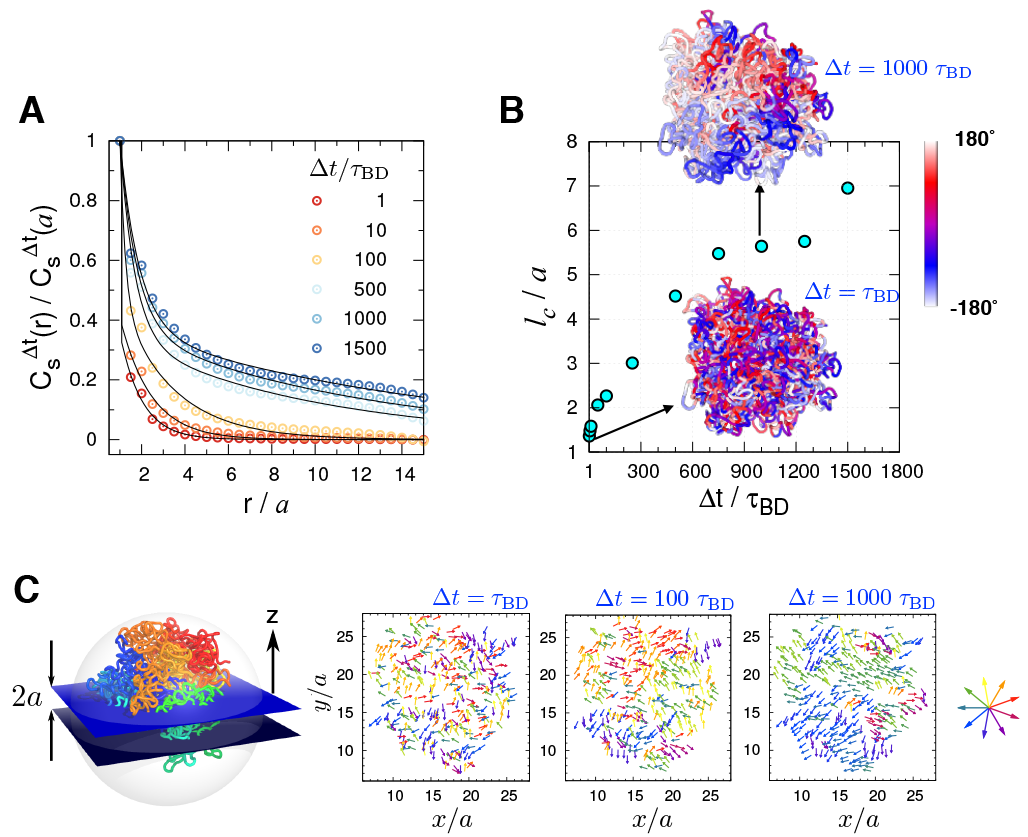
Spatial correlation between loci displacements. (A) Spatial correlation of loci displacements 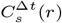 (Eq.4) with varying waiting time (Δ*t*). (B) Correlation length 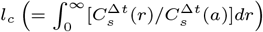 as a function of Δ*t*. Visualized on 3D chromosome structure are the displacement correlations of chromatin loci probed at short and large time gap (Δ*t* = *τ*_BD_ and 10^3^ *τ*_BD_) projected onto the xy-plane. The color-code on the structures depicts the azimuthal angle of loci displacement. (C) The displacement vector of loci in the equator plane are color-coded by direction. In each panel, the displacement vectors 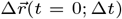 are calculated for Δ*t* = 1, 100, 1000 *τ*_BD_. Direction-dependent color scheme is shown on the right.

### Scale-dependent chromatin relaxation time

We explored the dynamical stability of chromosome structure at varying length scales. We calculated the time-evolution of the averaged mean square deviation of the distances between two loci with respect to the initial value (see Fig. 5A and the caption for the definition of *δ*(*t*)). Within our simulation time (*τ*max = 4 × 10^4^ *τ*_BD_), the largest value *δ*_max_(= 4.0 ± 0.3 *a*) is smaller than the value, *δ_c_* = 4.5 *a*, chosen to define different conformational clusters in Fig.1B. An extrapolation of *δ*(*t*) to *δ*(*τ_c_*) = *δ_c_* gives an estimate of *τ_c_* ≈ 10^5^ *τ*_BD_ ≈ 1.4 hours, which is a long time scale considering that most cells of adult mammals spend about 20 hours in the interphase (42).

**Fig. 5.**
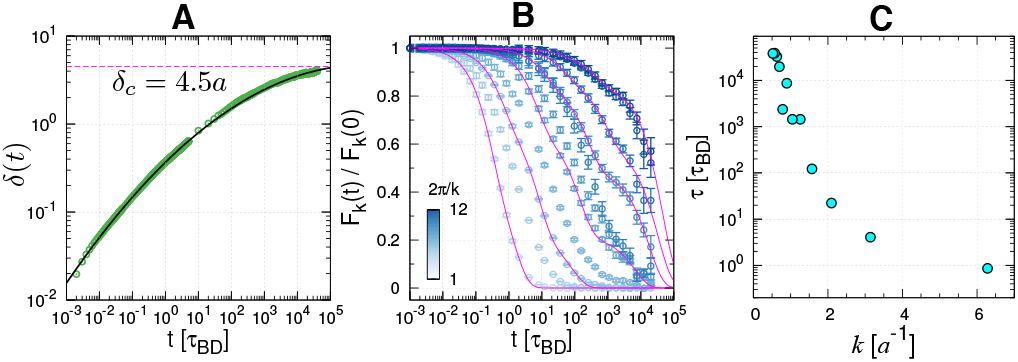
Lifetimes of chain conformations. (A) Time evolution of the root mean square distance between a pair of loci *r_i,j_* (*t*) at time *t* relative to its initial value (*r_i,j_* (0)) averaged over all pairs, defined by 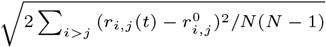. (B) Normalized intermediate scattering function *F_k_* (*t*)/*F_k_* (0), with different values of wave number *k*, calculated from BD simulation trajectories of the chromosome. (C) The chain relaxation time (*t*) for different wave number *k* is estimated by evaluating 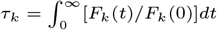.

From the definition of, *δ*(*t*), it follows that lim_*t*→∞_〈*δ*(*t*)〉 = *δ*_eq_. Here, *δ*_eq_ is finite, and 〈…〉) is an ensemble average, meaningful only if equilibrium is reached. The finiteness of *δ*_eq_ is a consequence of the polymer nature of the chromosomes. We estimate *δ*_eq_ assuming that the long time limit of the mean deviation of the distance between two loci is approximately the mean end-to-end distance between the loci. Thus, 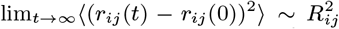 where *R_ij_* is the mean end-to-end distance between *i^th^* and *j^th^* loci. For |*i* − *j*| ≫ 1, we expect that 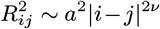. Consequently, *δ*_eq_ can be calculated using 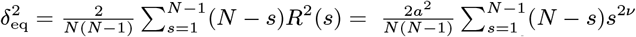. For *N* = 2712, and with *ν* = 1/3 we estimate *δ*_eq_ ≈ 9.4 *a*, which is greater than the value (*δ*_max_ ≈ 4.0 *a*) reached at the longest times in the simulations (Fig. 5A). An upper bound for *δ*_eq_ is 16.4 *a* (see SI). These considerations suggest that the chromosome dynamics is far from equilibrium on the time scale of a single cell cycle.

The scale-dependent relaxation dynamics of the chromatin domain is quantified using the time evolution of intermediate scattering function *F_k_* (*t*) (43, 44) calculated at different length scale (~ 2π/*k*) (Fig. 5B).

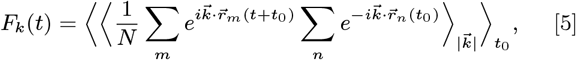

where 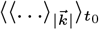 is an average over *t*_0_ and over the direction of vectors 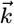 with magnitude 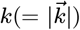. *F_k_*(*t*) shows that the chromatin chains are locally fluid-like (2π/*k* ≲ *a*), which is reminiscent of the recent analysis on the structural deformation of TADs (8), but their spatial organizations on intermediate to global scales (2π/*k* ≫ *a*) are characterized by slow relaxation dynamics. This scale-dependent relaxation time is reminiscent of a similar finding in random heteropolymers (45).

Relaxation time (*τ*) of a subdomain, whose size is *ξ* = 2π/*k*, can be estimated by evaluating 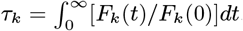, which can in turn be related to the number of segments comprising the subdomain as *ξ* ~ 2π/*k* ~ *s*^1/3^. Since the chromosome domain loses memory of the initial conformation by diffusion, the relaxation time *τ* is expected to obey *τ* ~ *ξ*^2^/*D*_eff_ ~ (*s*^1/3^)^2^/(*D*_0_/*s*) ~ *s*^5/3^. The relaxation times estimated from our chromosome model indeed scales with the domain size as *τ* ~ *s*^5/3^ (cyan symbols and solid line in Fig.6C).

**Fig. 6.**
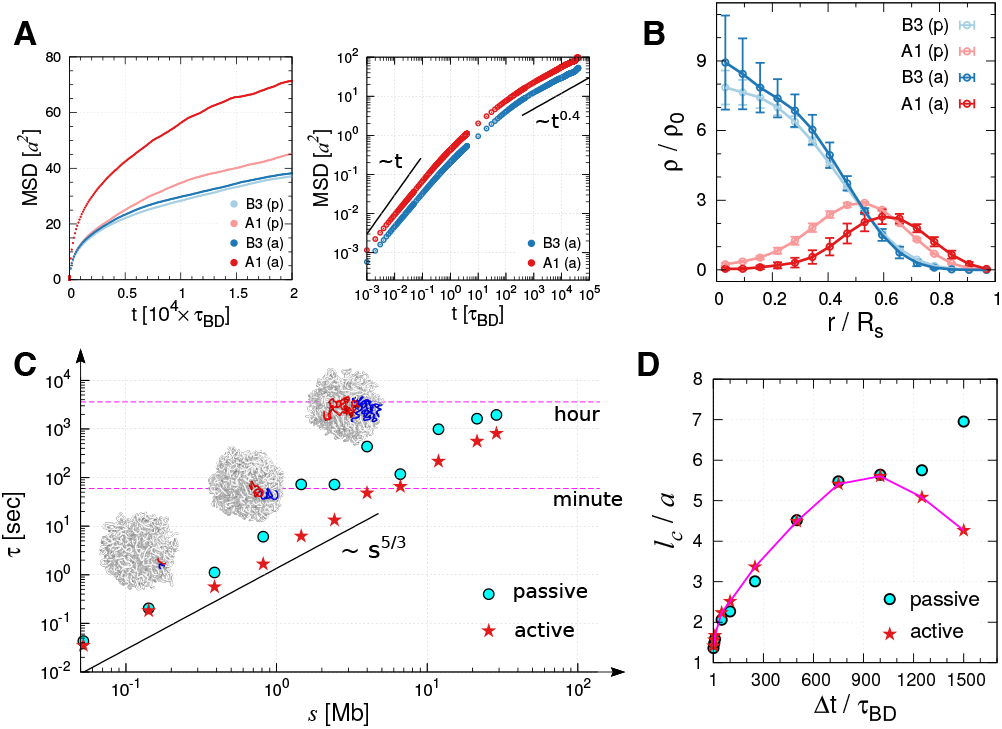
Effects of active force on chromosome organization and dynamics. (A) MSD of active and inactive loci compared with those under passive condition. Log-log plot is shown in the panel on the right. (B) Distribution of active (A1) and inactive (B3) loci with (a) and without (p) active force. In the presence of active force, the segregation of active and inactive loci is more evident. (C) Relaxation times estimated from the intermediate scattering functions. The wave number *k* was mapped to the corresponding number of loci inside the volume defined by the wave number. The red star symbols, the relaxation times in the presence of active forces, are depicted for the comparison with those under passive condition. (D) Correlation lengths for varying *Δt* calculated using the loci displacement correlations under passive (Fig.4A) and active (Fig. S4) conditions are compared.

### Effects of active forces on chromosome dynamics

Thus far, the findings from our simulations are based on using only passive forces in dictating chromatin dynamics. It could be argued that such a model neglects the most critical component of living systems. Live cells abound in a plethora of activities such as replication, transcription, and error-correcting dynamics. While these processes produce local directionality, when mapped onto the phenomenological description, the effects of vectorial forces on the surrounding environment at time scale longer than the correlation time of active noises can be assumed isotropic. We study how an increased noise strength 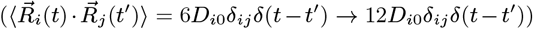 (46, 47) on the A1 and A2 monomers occupying 40 % of loci population for Chr10, which are classified as the active loci based on the epigenetic information (3), affects the dynamical properties of entire chromosome.

In the presence of active forces, while the diffusion exponent (*β* in MSD~ *t^β^*) is unaltered, the average MSD of A1 loci exhibits ~ 70 % increase relative to the passive case (Fig. 6A). The disproportionate increase in the mobility of A and B type monomers promotes the phase segregation of the two monomer types (Fig.6B, and see SI Movies 1 and 2). The active forces push A-type monomers towards the surface of the chromosome, and B-type monomers are pulled towards the center to offset this effect.

In terms of Fourier modes, the active forces mainly influence the chain relaxation described by the low frequency modes. For the high frequency modes or at local length scales (*k* ≳ 2π/3*a*), the intermediate scattering function is practically indistinguishable between active and passive cases (Fig. S3). The chromatin domains in the presence of active forces, on average, relax faster when the domain size is greater than the sub-Mb. A comparison of the relaxation times in Fig.6C under passive and active conditions highlight this difference.

Similarly, the effect of active forces on the correlation length (*l_c_*) is evident only at large waiting time (Δ*t*). We find that (*l_c_*) increases with Δ*t* under the passive condition, whereas a decrease of *l_c_* is observed for large Δ*t* under active force (Fig.6D). There is no distinction between the effects of passive and active forces on *l_c_* for small Δt; however, they deviate from each other for Δ*t* > 10^3^*τ*_BD_ ≈ 50 sec (Fig.6D). It is noteworthy that a similar dependence of correlation length with Δ*t* has been discussed in DCS measurement on genome-wide dynamics of live cell (9). Compared to thermal noise, active noise randomizes the global structure of chromatin chain more efficiently, which shortens the correlation length at sufficiently large lag time.

## Conclusions

Our study highlights the importance of chromosome architecture in determining the subdiffusive behavior and dynamic correlations between distinct loci. Most notably, we have shown that *structure* alone explains many of the dynamical features observed in living cells (9). In other words, chromosome organization dictates its dynamics. Remarkably, several static and dynamic properties of the model, including 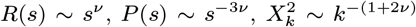, MSD~ *t*^2*ν*/(2*ν*+1)^, and *τ* ~ *s*^2*ν*+1^, are fully explained by the SF organization characterized by the single universal Flory exponent *ν* = 1/3, offering a unified perspective on both the structure and dynamics of chromosomes.

The relaxation time (*τ*) of the chromatin domain spans several orders of magnitude depending on its size (*s*), satisfying the scaling relation *τ* ~ *s*^5/3^ (Fig.6C). To be more concrete, while local chromatin domains of size *s* ≲ Mb, which include TADs and subcompartments, continuously reorganize on the time scale of *t* < 10^3^*τ*_BD_ ~ 50 seconds, it takes more than hours to a day for an entire chromosome chain (≳ 100 Mb) to lose memory of its initial conformation. This timescale associated is expected to grow even further at higher volume fractions (22). It is likely that under *in vivo* conditions, with 46 chromosomes segregated into chromosome territories, the time scale for relaxation can be considerable.

The effects of active forces on chromatin dynamics (9, 48) deserve further discussion. While active forces enhance chain fluctuations and structural reorganization, the effect on chromatin domain manifests itself only on length scales greater than 5.5 *a* (≈ 0.8 *μm*), and on a time scale greater than 50 sec (Fig.6D). This is closely related to the active cytoskeletal network using microrheology measurements (49), where the effect of myosin activity is observable only at low frequencies in the power spectrum of the response function. Of course, the active forces in live cell nuclei is not a scalar, and it remains a challenge to model their vectorial nature in the form of force dipole or vector force in the context of chromatin dynamics (46). Vector activities would render loci with super-diffusive motion (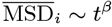 with 1 < *β* < 2) dominant, and could in principle elicit a qualitative change in the dynamical scaling relations. However, the dynamic scalings discussed in this study (e.g., MSD~ *t*^0.4^) are in good agreement with those observed in interphase chromatins (9, 12, 34). In terms of *power* generated in a cell, the passive (thermal) power *W_p_* ~ *k_B_T*/ps is many orders of magnitude greater than the active power (e.g., molecular motors, *W_a_* ~ 20 *k_B_T*/10 ms (42)). At least in the interphase, the gap between the total passive and active power is substantial, because the number of active loci (*N_a_*) is smaller than the number of passive loci (*N_p_*), satisfying the relation *N_p_ W_p_* ≫ *N_a_W_a_*. The robustness of the diffusion exponent indicates that the total contribution of the scalar and vector activities during the interphase is negligible compared to thermal agitation, and does not entirely offset the effects of chromosome architecture on the dynamics.

Taken together, our study unequivocally shows that chromosome architecture alone, captured by the single Flory exponent, determines much of the loci dynamics during the interphase.

## Materials and Methods

To build the chromosome 10 model of human lymphoblastoid cell, we employed the potential in MiChroM. The coarse graining of chromatin leads to *N* = 2712 loci with the diameter of each *a* ≈ 150 nm. Thus, 50 Kb of DNA is in a single locus. The inverse mapping of the Hi-C map to the ensemble of chromosome structures was carried out by sampling the conformational space using low-friction Langevin simulations (31). The generated structures follow the characteristic scaling of the contact probability, *P*(*s*) ~ *s*^−1^, and reproduce the spatial distribution of A/B compartment as well as the plaid pattern noted in Hi-C experiments. To study the dynamics of chromatin, we used Brownian dynamics. The Brownian time tbd ≈ 50 ms in physical time. The details of the energy function and simulation algorithm are provided in the SI.

## Acknowledgments

We thank the Center for Advanced Computation in KIAS for providing computing resources. D.T. acknowledges support from the National Science Foundation (CHE 16-32756 and 16-36424) and the Collie-Welch Chair (F-0019).

## References

1. Lieberman-Aiden E, et al.(2009) Comprehensive mapping of long-range interactions reveals folding principles of the human genome. Science 326(5950): 289–293.

2. Dixon JR, et al.(2012) Topological domains in mammalian genomes identified by analysis of chromatin interactions. Nature 485(7398): 376–380.

3. Rao SSP, et al.(2014) A 3D Map of the Human Genome at Kilobase Resolution Reveals Principles of Chromatin Looping. Cell 159(7): 1665–1680.

4. Du Z, et al.(2017) Allelic reprogramming of 3D chromatin architecture during early mammalian development. Nature 547(7662): 232–235.

5. Shen Y, et al.(2012) A map of the cis-regulatory sequences in the mouse genome. Nature 488(7409): 116–120.

6. Sanyal A, Lajoie BR, Jain G, Dekker J (2012) The long-range interaction landscape of gene promoters. Nature 489(7414): 109–113.

7. Stevens TJ, et al.(2017) 3D structures of individual mammalian genomes studied by singlecell Hi-C. Nature 544(7648): 59–64.

8. Tiana G, et al.(2016) Structural fluctuations of the chromatin fiber within topologically associating domains. Biophys. J.110(6): 1234–1245.

9. Zidovska A, Weitz DA, Mitchison TJ (2013) Micron-scale coherence in interphase chromatin dynamics. Proc. Natl. Acad. Sci. USA 110(39): 15555–15560.

10. Hajjoul H, et al.(2013) High-throughput chromatin motion tracking in living yeast reveals the flexibility of the fiber throughout the genome. Genome Res. 23(11): 1829–1838.

11. Lucas JS, Zhang Y, Dudko OK, Murre C (2014) 3D Trajectories Adopted by Coding and Regulatory DNA Elements: First-Passage Times for Genomic Interactions. Cell 158(2): 339–352.

12. Bronshtein I, et al.(2015) Loss of lamin A function increases chromatin dynamics in the nuclear interior. Nat. Commun.6: 8044.

13. Cremer T, et al.(2015) The 4D nucleome: Evidence for a dynamic nuclear landscape based on co-aligned active and inactive nuclear compartments. FEBS Lett. 589(20 Pt A): 2931–2943.

14. Nagano T, et al.(2017) Cell-cycle dynamics of chromosomal organization at single-cell resolution. Nature 547(7661): 61–67.

15. Dekker J, et al.(2017) The 4d nucleome project. Nature 549(7671): 103499.

16. Sanborn AL, et al.(2015) Chromatin extrusion explains key features of loop and domain formation in wild-type and engineered genomes. Proc. Natl. Acad. Sci. USA 112(47): E6456–E6465.

17. Fudenberg G, et al.(2016) Formation of Chromosomal Domains by Loop Extrusion. Cell Reports 15(9): 2038–2049.

18. Grosberg, NechaevSK, Shakhnovich EI (1988) The role of topological constraints in the kinetics of collapse of macromolecules. J. Phys.49(12): 2095–2100.

19. Mirny LA (2011) The fractal globule as a model of chromatin architecture in the cell. Chromosome Res. 19(1): 37–51.

20. Halverson JD, Smrek J, Kremer K, Grosberg AY (2014) From a melt of rings to chromosome territories: the role of topological constraints in genome folding. Rep. Prog. Phys. 77(2): 022601.

21. Bohn M, Heermann DW, van Driel R (2007) Random loop model for long polymers. Phys. Rev. E 76: 051805.

22. Kang H, Yoon YG, Thirumalai D, Hyeon C (2015) Confinement-Induced Glassy Dynamics in a Model for Chromosome Organization. Phys. Rev. Lett.115: 198102.

23. Wang S, Xu J, Zeng J (2015) Inferential modeling of 3d chromatin structure. Nucleic Acids Res. 43(8): e54.

24. Szalaj P, et al.(2016) 3D-GNOME: an integrated web service for structural modeling of the 3D genome. Nucleic Acids Res. 44(W1): W288.

25. Tjong H, et al.(2016) Population-based 3d genome structure analysis reveals driving forces in spatial genome organization. Proc. Natl. Acad. Sci. USA 113(12): E1663–E1672.

26. Di Stefano M, Paulsen J, Lien TG, Hovig E, Micheletti C (2016) Hi-C-constrained physical models of human chromosomes recover functionally-related properties of genome organization. Sci. Rep.6: 35985.

27. Di Pierro M, Zhang B, Aiden EL, Wolynes PG, Onuchic JN (2016) Transferable model for chromosome architecture. Proc. Natl. Acad. Sci. USA 113(43): 12168–12173.

28. Zhang B, Wolynes PG (2015) Topology, structures, and energy landscapes of human chromosomes. Proc. Natl. Acad. Sci. USA 112(19): 6062–6067.

29. Nagano T, et al.(2013) Single-cell Hi-C reveals cell-to-cell variability in chromosome structure. Nature 502(7469): 59–64.

30. Ramani V, et al.(2017) Massively multiplex single-cell Hi-C. Nat. Methods 14(3): 263–266.

31. Honeycutt JD, Thirumalai D (1992) The nature of folded states of globular proteins. Biopolymers 32(6): 695–709.

32. Lua RC (2012) PyKnot: a PyMOL tool for the discovery and analysis of knots in proteins. Bioinformatics 28(15): 2069.

33. Mortazavi A, Williams BA, McCue K, Schaeffer L, Wold B (2008) Mapping and quantifying mammalian transcriptomes by RNA-Seq. Nat. Methods 5(7): 621–628.

34. Shinkai S, Nozaki T, Maeshima K, Togashi Y (2016) Dynamic Nucleosome Movement Provides Structural Information of Topological Chromatin Domains in Living Human Cells. PLOS Comput. Biol.12(10): 1–16.

35. Liu L, Hyeon C (2016) Contact statistics highlight distinct organizing principles of proteins and rna. Biophys. J.110(11): 2320–2327.

36. Shi G, Liu L, Hyeon C, Thirumalai D (2017) Interphase Human Chromosome Exhibits Out of Equilibrium Glassy Dynamics. bioRxiv.

37. Tamm MV, Nazarov LI, Gavrilov AA, Chertovich AV (2015) Anomalous Diffusion in Fractal Globules. Phys. Rev. Lett.114: 178102.

38. Weber SC, Theriot JA, Spakowitz AJ (2010) Subdiffusive motion of a polymer composed of subdiffusive monomers. Phys. Rev. E 82: 011913.

39. Lampo TJ, Kennard AS, Spakowitz AJ (2016) Physical Modeling of Dynamic Coupling between Chromosomal Loci. Biophys. J.110(2): 338–347.

40. Panja D (2010) Anomalous polymer dynamics is non-Markovian: memory effects and the generalized Langevin equation formulation. J. Stat. Mech. Theory E 2010(06): P06011.

41. Doi M, Edwards SF (1988) The Theory of Polymer Dynamics (International Series of Monographs on Physics). (Oxford University Press).

42. Milo R, Phillips R (2016) Cell biology by the numbers. (Garland Science, New York).

43. Hansen JP, McDonald IR (2006) Theory of Simple Liquids. (Academic Press), 3 edition.

44. Liu L, Pincus PA, Hyeon C (2017) Heterogeneous Morphology and Dynamics of Polyelectrolyte Brush Condensates in Trivalent Counterion Solution. Macromolecules 50(4): 1579–1588.

45. Thirumalai D, Ashwin V, Bhattacharjee JK (1996) Dynamics of random hydrophobic-hydrophilic copolymers with implications for protein folding. Phys. Rev. Lett.77(27): 5385–5388.

46. Bruinsma R, Grosberg AY, Rabin Y, Zidovska A (2014) Chromatin hydrodynamics. Biophys. J.106(9): 1871–1881.

47. Smrek J, Kremer K (2017) Small activity differences drive phase separation in active-passive polymer mixtures. Phys. Rev. Lett. 118(9): 098002.

48. Weber SC, Spakowitz AJ, Theriot JA (2012) Nonthermal ATP-dependent fluctuations contribute to the in vivo motion of chromosomal loci. Proc. Natl. Acad. Sci. USA 109(19): 7338–7343.

49. Mizuno D, Tardin C, Schmidt C, MacKintosh F (2007) Nonequilibrium mechanics of active cytoskeletal networks. Science 315(5810): 370–373.

